# The Energetics of Maintaining the Lateral Balance Are Terrain-Specific; a Normal Lookahead Significantly Reduces Active Balance Maintenance Work

**DOI:** 10.1101/2024.07.16.603705

**Authors:** Seyed-Saleh Hosseini-Yazdi

## Abstract

Humans must actively control their lateral balance through frontal plane work or adjusting lateral foot placement. With constant muscle efficiency and considering the energetic consequences of Center of Mass (COM) work variability, we estimated the metabolic cost of lateral balance maintenance and compared it with the Workman model. Like the Workman model, we found that lateral balance energetics were mainly associated with terrain amplitude. Increased walking speed’s effect on step transition work might be offset by reduced step width. A significant rise in lateral work magnitude (+157.1%) with restricted lookahead was potentially linked to wider steps. Comparing mechanical work with the Workman model, we found significant differences in magnitudes, suggesting that the Workman model included additional costs such as force rate generation, muscle coactivation, or posture maintenance not reflected in the COM lateral work and its variability.

**Practitioner Summary:** Lateral balance maintenance during walking requires active regulation, yet it is not studied mechanically. Our study found that lateral control costs are terrain-specific and increase with restricted lookahead. With age, only variability increases. We assume associated energetics rise with both work and variability.

## Introduction

Human walking is considered unstable in the Mediolateral (M/L) direction. Thus, humans need to modulate their balance actively [1]. This dynamic contradicts the human-proposed walking characteristic of the sagittal plane [2]. To maintain balance, humans must utilize active controls based on sensory inputs (vision, vestibular, etc.) [3]. Accordingly, the modulated foot placement is proposed as one approach that feeds on visual information [1]. Since the lateral foot placement must occur during the step-to-step transition at which the Center of Mass (COM) velocity is adjusted for the subsequent step [4], it is plausible to consider that in addition to the mechanical work exerted to energize the forward progression [5,6], humans also perform positive mechanical work to regulate COM motion and balance in the frontal plane [7]. Such work must exact a metabolic cost contributing to overall walking energetics. Since uneven walking may elevate stability maintenance challenges, the M/L work performance and ongoing corrections [8] may also alter. Therefore, variations in energy expenditure must occur. Walking velocity, age, and state of lookahead may also affect the performed lateral work and its associated energetics.

During uneven walking, induce perturbations affect the human walking pattern. It has been observed that over complex terrains, humans decrease their walking velocity [9], undermining the optimal relationship between walking velocity and step length [10]. This represents a trade-off involving energy economy, safety, balance control, stability, and other factors [9], wherein foot placement deviates from the typical pattern observed during even walking [10,11]. Uneven terrains also affect the walking periodicity in which one step is different from others. For instance, step elevation change may cause instantaneous velocity fluctuations [12] when other factors are kept the same. Such fluctuations may affect the balance considerations [8]. Elevated gait variability might be a consequence of deviation from periodicity with a greater need for additional active control [13], manifested through increased force or work exertion. This increased variability is also linked to elevated walking energetics [8].

A simple model of powered walking may explain the mechanics of transitioning between steps in both the sagittal and frontal planes [5,7]. Perturbations in the sagittal plane, such as elevation changes, occur in direction that exhibits passive stability, leading to decay and eventual restoration of stability [1,2]. Conversely, lateral motion is believed to possess inherent instability, which can be compensated by widening the step [7] or active control [1]. Visual cues also play a crucial role in stabilizing the COM laterally, assisting in selecting appropriate foot placement [14]. Given that lateral stability can be achieved with minimal effort through lateral foot placement [1], increasing step width serves as an indicator of the effort in maintaining stability and balance. Nevertheless, wider steps necessitate greater mechanical work to redirect the COM in the frontal plane [7]. The step-up and step-down characteristic of uneven walking may further change the mechanical work required to propel lateral movement (Figure 1).

**Figure 1:**
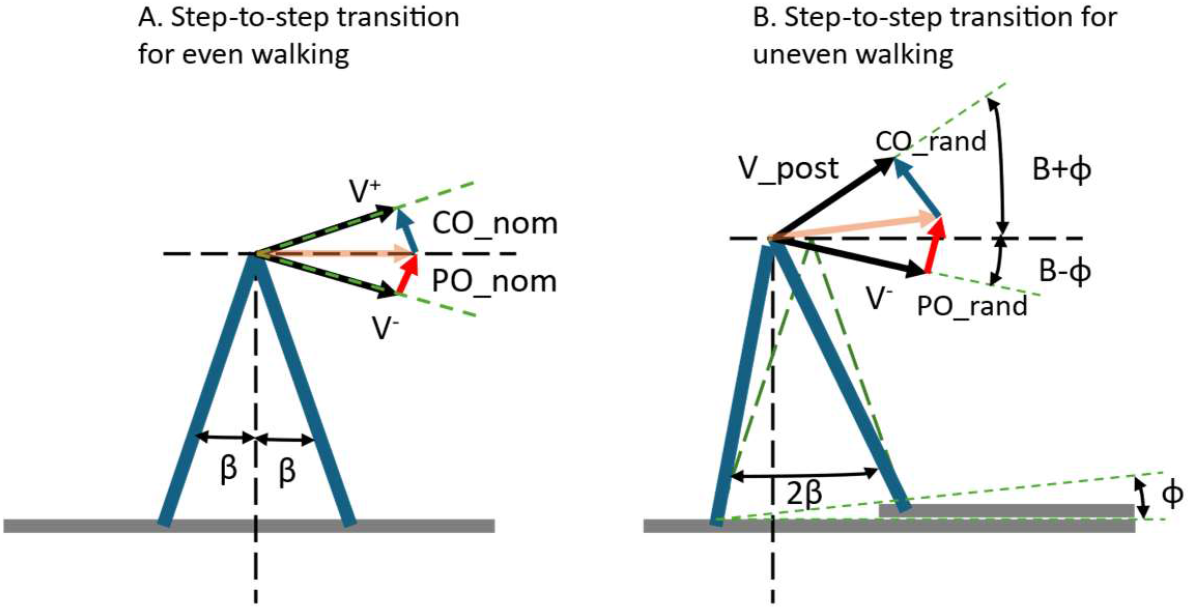
Schematics showing the step-to-step transition of the Center of Mass (COM) in the frontal plane. (A) Redirection of COM walking over even terrain. (B) The redirection of COM walking over an uneven terrain. The redirection happens with the exertion of push-off, which is followed by a collision. The push-off magnitude alters with step width (*β*) and walking velocity. For small angle approximation, the push-off estimate is *po* = *v*_ave_ · *β*.

The denial of visual information during uneven walking or the Central Nervous System (CNS) state, which is proposed to deteriorate with old age [15], may also make lateral balance control more challenging [16]. With the restricted lookahead, humans may not choose the proper foot placement that enables them to extract the maximum energy from the COM inverted pendulum motion at the point of transition [14]. Therefore, they may also need to exert additional positive work in the frontal plane.

It is proposed that balance cost may be estimated from the general walking energetics trajectories. Based on the even walking stable energetics features, Workman et al. [17] have suggested a three-compartment energetics model. The first compartment is associated with the basal metabolic rate, measured at rest conditions, which exhibits the human body’s metabolism rate. The second compartment is termed as the general metabolic of walking. They proposed that the general walking metabolic rate appears during walking and stays almost constant regardless of the walking velocity. Therefore, they assumed it to be unrelated to walking movement and mainly used to maintain balance and posture. The third compartment is deemed to have the metabolic rate associated with walking movement as it increases with walking velocity.

Given the proposition that COM mechanical work can explain the energetic cost of walking [6], an examination of positive lateral COM work during uneven walking not only explain changes in work magnitude but also potentially signifies associated energetic variations influenced by terrains and other conditions. Consequently, we hypothesize that during uneven walking, particularly with the restricted lookahead, the mechanical work required to sustain balance and modulate ongoing movements increases. Therefore, we expect the associated energetic cost also to escalate. Here, we aim to evaluate if a proposed simple model for balance energetics can be supported by a mechanistic approach.

## Materials and Methods

We adjusted an instrumented treadmill structure (Bertec Corporation, Columbus, Ohio, USA) for our uneven walking experiments. This customization enabled us to replicate the effects of uneven walking by mounting uneven terrains onto the extended support structure of the treadmill’s moving section [18]. Fabricating these terrains involved attaching construction foam blocks to the belts of conventional treadmills with varying heights (Max_h = 0.019 m, 0.032 m, and 0.045 m). These foam blocks, securely affixed with construction adhesive, spanned the width of the belt. We maintained a consistent 0.3 m height for each elevation, allowing for either level landing or a slight incline spanning two adjacent heights.

We also created an uneven terrain by applying the construction glue to the surface of the belt to create a constant contact with the terrain irregularities (Max_h = 0.005 m, Figure 2A). These terrains were designed to be open-ended, allowing them to wrap around the instrumented treadmill’s original belt when the tensioning rollers were released. Closing the terrains with alligator lacing clips, the tensioning rollers were reset to their original positions.

**Figure 2:**
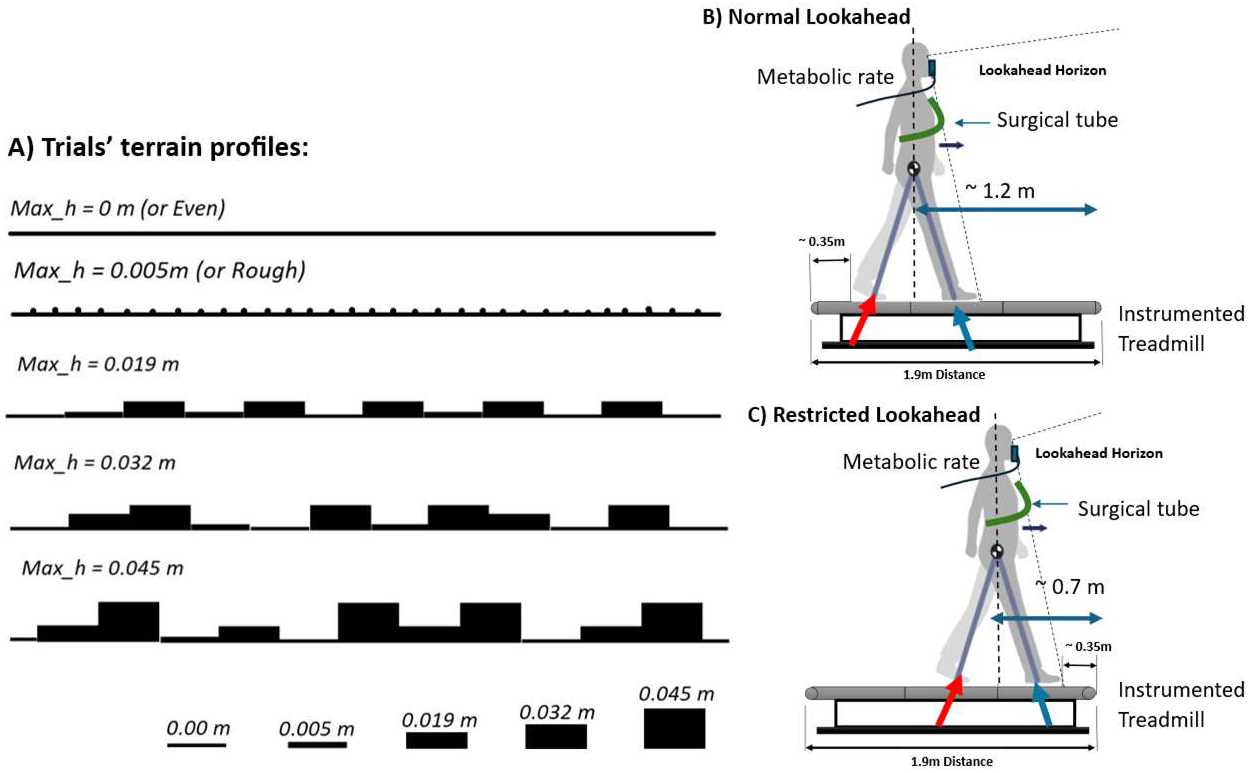
(A) the profiles of the uneven terrains used for the experiments. They were wrapped around the existing belt and tensioned simultaneously. (B) walking with normal lookahead in which the subjects had about two step-lengths view of the coming terrain. A surgical tube mounted on a tripod on either side of the treadmill was used to assist subjects in keeping their relative positions constantly. (C) walking with the restricted lookahead in which subjects view was limited.

We invited two groups of young adults (age: 27.69 ± 5.21, mass: 71.0 ± 6.16 kg, six females and four males) and older adults (age: 66.1 ± 1.44, mass: 77.34 ± 12.13 kg, six males and four females) to participate in our experiment. Each subject provided written informed consent before engaging in the experiments. The experimental procedures received approval from the University of Calgary Board of Ethics.

The uneven walking trials were conducted under Normal and Restricted lookaheads. In the Restricted Lookahead, subjects were positioned approximately 0.7 meters from the front end of the treadmill. We encouraged them to walk naturally while their view of the front was limited. Conversely, subjects were placed 1.2 meters behind the front end in the Normal Lookahead, providing two-step horizon visibility of approaching terrain (Figure 2B & C). A flexible surgical tube was installed across the treadmill on two tripods to facilitate subjects maintaining their positions. Subjects were instructed to contact the tube gently with their torsos, and verbal feedback was provided when necessary. Trial orders were randomized (Table 1).

**Table 1:** Trials schedule of uneven walking. It shows the walking velocity (*m* · *s*^-1^) and terrain amplitude (m) used for each age group. YA: young adults, OA: older adults. Upper-case letters indicate trials with normal lookahead, and lower-case letters refer to restricted lookahead trials.

| Terrains Max_h (m) | 0 | Rough | 0.019 | 0.032 | 0.045 |
| --- | --- | --- | --- | --- | --- |
| Trials Velocities (m/s) |  |  |  |  |  |
| 0.8 | YA | YA | YA | ya-oa/YA-OA | YA |
| 1 | YA | YA | YA | ya-oa/YA-OA | YA |
| 1.2 | ya-oa/YA-OA | YA | ya-oa/YA-OA | ya-oa/YA-OA | ya-oa/YA-OA |
| 1.4 | YA | YA | YA | ya-oa/YA-OA | YA |

Each trial ran for six minutes. We analyzed metabolic cost from respirometry data, averaging oxygen consumption rates and carbon dioxide production for the last three minutes of each six-minute trial (Cosmed K5 metabolic system, Cosmed S.r.l., Rome, Italy). Net Metabolic Rates (NMR) were derived by deducting each subject’s average standing metabolic rate. NMRs were normalized using subject mass (M) to facilitate comparison across subjects of different sizes.

We estimated the energetics of balance and posture following the Workman model [17]. To calculate the energetics of the second compartment, we fitted a third-degree polynomial to the net metabolic rates of each terrain (^Ė^ ∼v^3^). The estimated intercepts were considered as the second compartment of walking energetics. Additionally, as it is proposed that the walking energetics increases with the square of the terrain (^Ė^ _balance_∼Max_h^2^) [19] we also fitted a second degree polynomial to the balance energetics.

The ground reaction forces (GRF) were collected at a 960 Hz sampling rate for sixty seconds. The gait events were determined based on the GRF data. The lateral COM velocity was calculated by time integration of lateral ground reaction forces (GRF) utilizing the method described by Donelan et al. [20]. We also calculated the M/L power by the dot product of COM lateral velocity and M/L forces [21]. The positive lateral COM power was integrated over the stance time to derive the active work. Then, the average lateral work rate was estimated by dividing the average lateral work by its associated average step duration [22]. Accordingly, the metabolic cost of lateral work was calculated, assuming a constant muscle efficiency for active work (25%) [22]. The influences of variabilities were applied, assuming even walking with normal lookahead for young adults (control) as the base case and calculating the energetics multiplier for raised variability based on O’Conner et al. [23].

For statistical analysis, we employed linear regression with Linear Mixed Effects Models (Statsmodels 0.15.0), where the dependent variable was the M/L work or its variability. Independent variables included trial walking velocities, terrain amplitudes (Max_h), age group (young adults (YA) versus older adults (OA)), and state of lookahead (normal lookahead versus restricted lookahead). Regression coefficients and associated *p*-values were reported to illustrate the relationships and significance between the variables of interest and the independent factors.

## Results

We quantified the M/L COM work and its variabilities to estimate the metabolic cost associated with maintaining balance and compared it with the estimate from the Workman model derived based on walking energetics. We could detect subtle changes with walking speed or terrain amplitude rise in lateral COM power trajectories. Nevertheless, the largest variation appeared to happen with the state of lookahead change (Figure 3).

**Figure 3:**
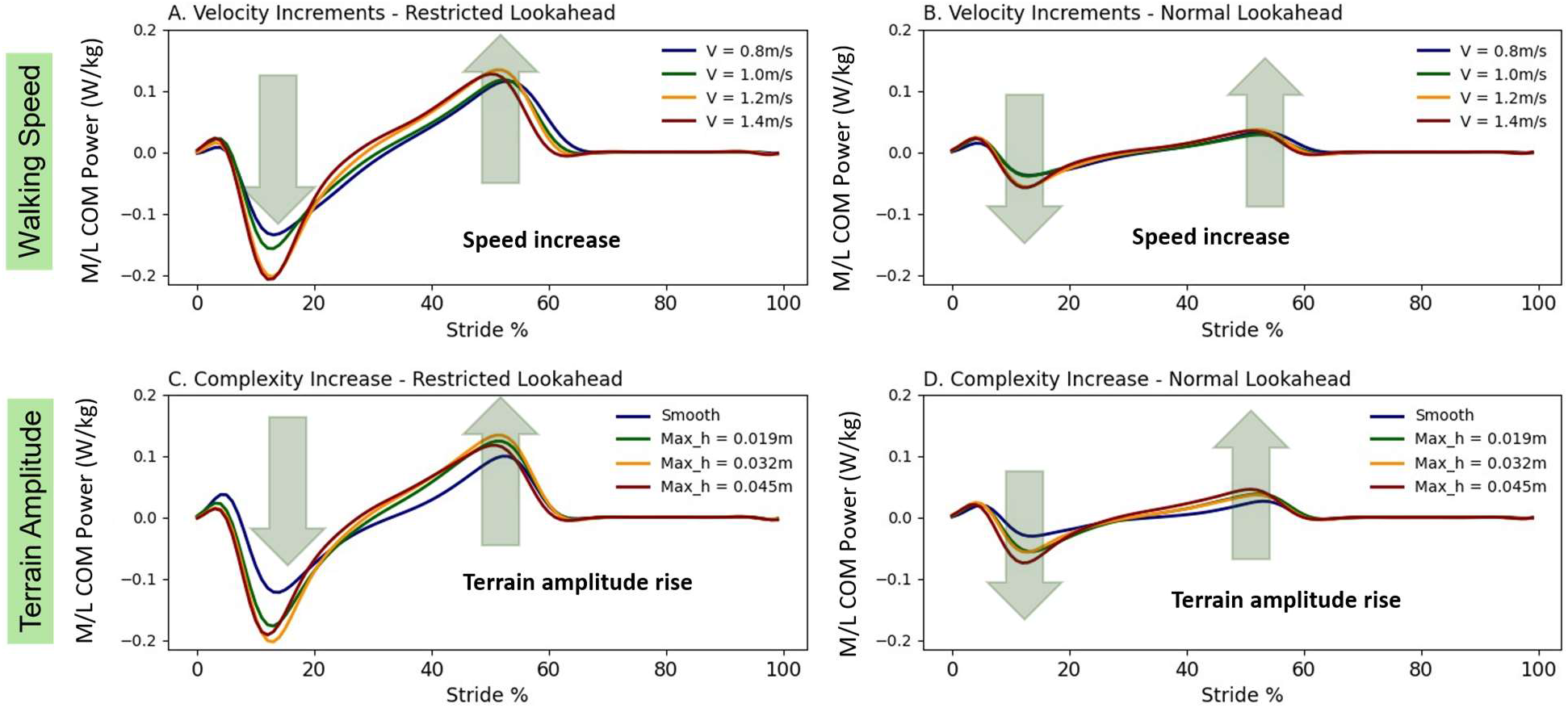
The lateral power trajectories with (top row: A. and B.) across walking speed with normal and restricted lookahead, and (lower row: C. and D.) across terrain amplitudes with normal and restricted lookahead. While the lateral COM power trajectories changes with terrain or walking speed were subtle, their peak appeared to change substantially with the restricted lookahead.

The M/L work changed significantly with terrain amplitude and state of lookahead. With the normal lookahead, M/L work rose from 0.010 *J* · *kg*^-1^ to 0.018 *J* · *kg*^-1^ (80%) across terrains (v = 1.2 *m* · *s*^-1^). Alternatively, with the restricted lookahead, the M/L work ranged from 0.032 *J* · *kg*^-1^ to 0.040 *J* · *kg*^-1^ (25%).

The M/L variability changed significantly with all the control parameters. The M/L increased with age while it reduced with normal lookahead (offsets). The variability ranges for the set categories were; young adults with normal lookahead: 0.004 *J* · *kg*^-1^ to 0.011 *J* · *kg*^-1^ (+175.0), young adults with restricted lookahead: 0.013 *J* · *kg*^-1^ to 0.020 *J* · *kg*^-1^ (+53.8%), older adults with normal lookahead: 0.007 *J* · *kg*^-1^ to 0.014 *J* · *kg*^-1^ (+100%), and older adults with restricted lookahead: 0.016 *J* · *kg*^-1^ to 0.023 *J* · *kg*^-1^ (43.8%), respectively (Table 2). The work rate and their variabilities were derived by dividing the M/L works with associated step durations (Figure 4A & B) Assuming a constant muscle efficiency [22] and the energetics influence of the gait variabilities (control: even walking for young adults with the normal lookahead) [8], we estimated net metabolic rate trajectories associated with the lateral work. With the normal lookahead, the young adults’ estimated M/L energetic rate rose from 0.072 W · kg^-1^ to 0.169 W · kg^-1^ (+134.7%), while for the older adults, it rose from 0.078 W · kg^-1^ to 0.180 W · kg^-1^(+130.8%). Likewise, with the restricted lookahead, the estimated energetics varied from 0.304 W · kg^-1^ to 0.454 W · kg^-1^ (+49.3%) and from 0.322 W · kg^-1^ to 0.478 W · kg^-1^ (+48.4%) for young and older adults respectively (Figure 4C).

**Table 2:**
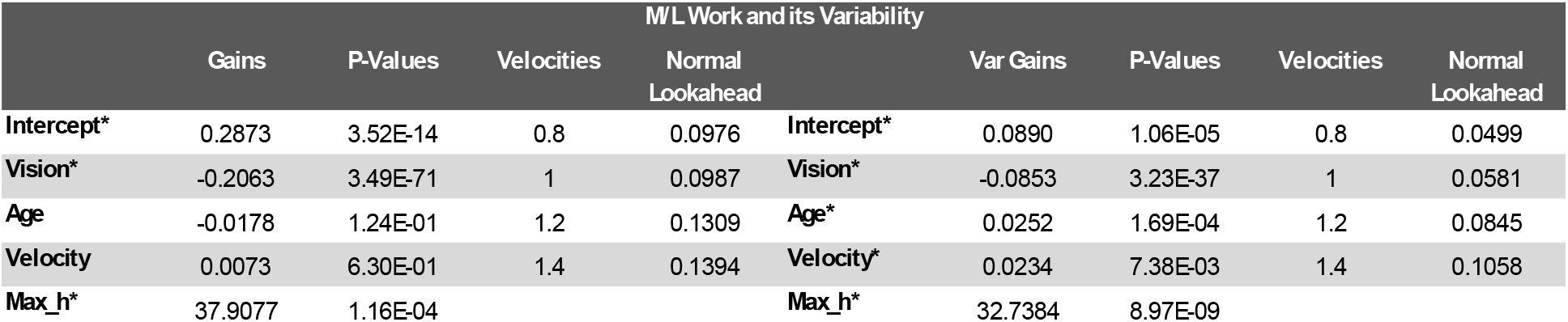
The statistical analysis results summary (linear mixed model) for COM lateral work and its variability. The young adults’ average lateral COM works and variabilities for normal lookahead are provided as a reference when subjects walked on a constant terrain (Max_h = 0.032 m) with different velocities. The significant variations are indicated by asterisks (*).

| M/L Work and its Variability |  |  |  |  |  |  |  |  |  |
| --- | --- | --- | --- | --- | --- | --- | --- | --- | --- |
|  | Gains | P-Values | Velocities | Normal Lookahead |  | Var Gains | P-Values | Velocities | Normal Lookahead |
| **Intercept\*** | 0.2873 | 3.52E-14 | 0.8 | 0.0976 | **Intercept\*** | 0.0890 | 1.06E-05 | 0.8 | 0.0499 |
| **Vision\*** | -0.2063 | 3.49E-71 | 1 | 0.0987 | **Vision\*** | -0.0853 | 3.23E-37 | 1 | 0.0581 |
| **Age** | -0.0178 | 1.24E-01 | 1.2 | 0.1309 | **Age\*** | 0.0252 | 1.69E-04 | 1.2 | 0.0845 |
| **Velocity** | 0.0073 | 6.30E-01 | 1.4 | 0.1394 | **Velocity\*** | 0.0234 | 7.38E-03 | 1.4 | 0.1058 |
| **Max_h\*** | 37.9077 | 1.16E-04 |  |  | **Max_h\*** | 32.7384 | 8.97E-09 |  |  |

**Figure 4:**
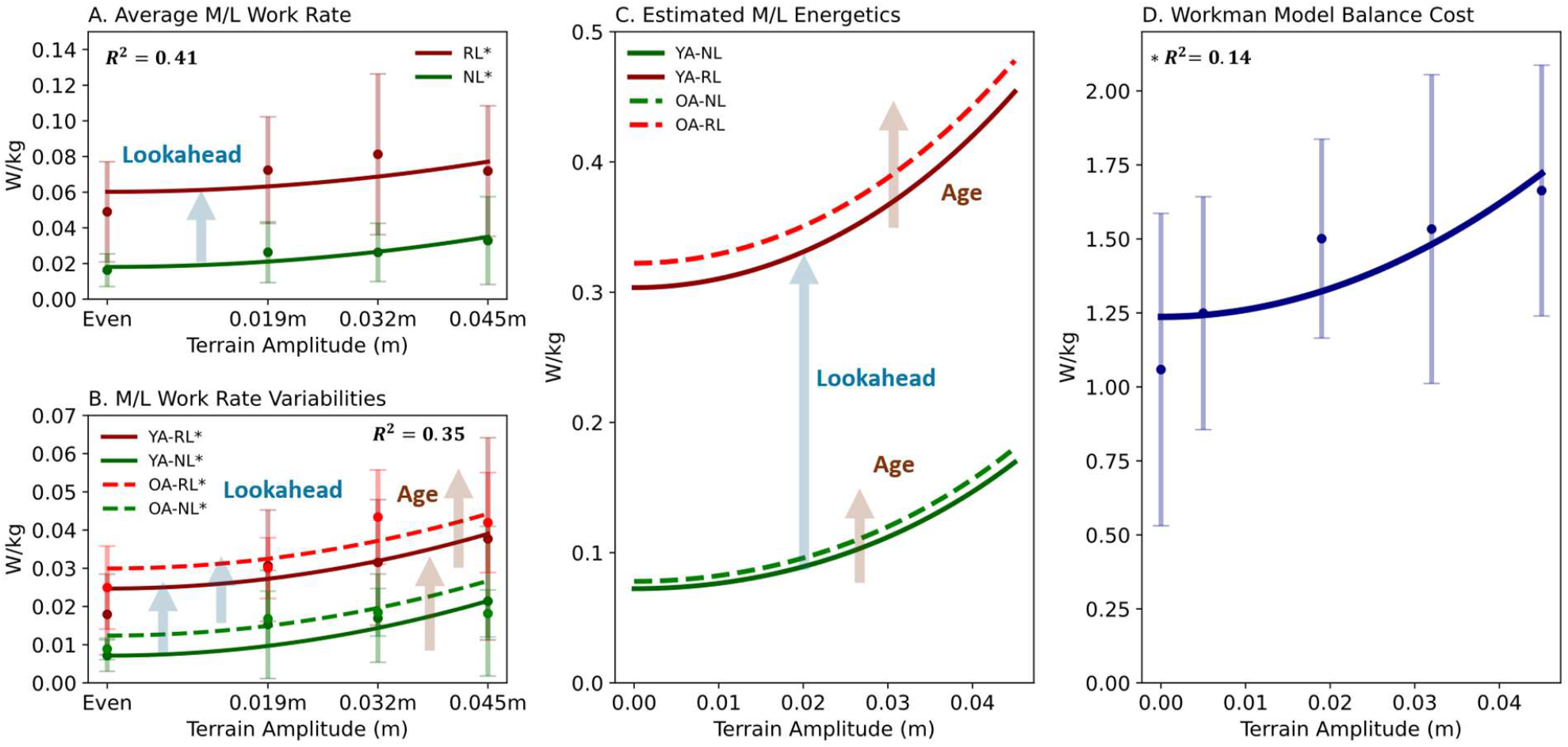
(A) The estimated ML work rate with terrain amplitude increase (B) The estimated ML work rate variability with the terrain amplitude rise (C) The estimated energetics of maintaining lateral balance assuming a constant muscle efficiency for active works (*η* = 0.25) and constant influence of variabilities (control: even walking for young adults with normal lookahead variability). The estimated lateral work energetics were only related to terrain amplitudes. (D) The Workman model prediction for lateral balance and posture maintenance: The Workman estimate magnitudes were more considerable than the COM lateral work estimates. It also suggested that the energetics of balance was only related to the terrains.

The Workman model’s balance and posture maintenance cost increased from 1.24 W · kg^-1^ to 1.72 W · kg^-1^ (+38.7%) across terrains (Figure 4D). At Max_h = 0.032 m, the cost of balance and posture rose from 1.53 W · kg^-1^ to 1.88 W · kg^-1^ for young adults with restricted lookahead (+22.9%), to 2.31 W · kg^-1^ for older adults with normal lookahead (+51.0%), and to 3.04 W · kg^-1^ for older adults with restricted lookahead (+98.7%), respectively.

## Discussion

It is shown that lateral balance maintenance is challenging and requires active regulations for which visual information plays a crucial role [1]. The regulation of the lateral balance in part may be done by the active muscle work performance [7]. Since the COM work roughly describes the walking energetics [20], the lateral mechanical work also may represent the balance energetic cost in walking.

We could show that the COM lateral work increases with the restricted lookahead substantially. As step width also increases with the vision impairment [24,25], it might explain the rise in the lateral work partially [5,7]. Therefore, the increased lateral work with the terrain amplitude increase may be also associated with the elevated step width over complex terrains [18]. A portion of increased COM lateral work increase with the restricted lookahead perhaps is also associated with the absence of proper foot placement in which maximum possible potential energy is converted into the kinetic energy [14]. Hence, the suboptimal foot placement demands a larger transition mechanical work, particularly over uneven terrains.

The sharp rise in COM lateral power peak and the associated work suggests that other senses contributions to the lateral balance control are far inferior to vision [3]. It additionally suggests that individuals considered vulnerable (e.g., older adults or otherwise with physical deficits) might rely more on visual cues to negotiate complex terrains [26]. Additionally, although relatively small but significant energetics expenditure rise associated with the lateral work might also contribute to exhaustion [27], loss of momentum, and possibly fall in vulnerable individuals [28].

We could not recognize any significant changes in the lateral work with walking speed increase or age. It seems that the step width reduction shown in higher walking speed [29] cancels the influence of increased speed in the step-to-step transition in the frontal plane (*po*_frontal_ = *v*_ave_ · tan *β*, Figure 1). Therefore, it is plausible to infer that the lateral balance is terrain specific (Figure 4C). However, if the step width is reduced sufficiently, it causes leg circumduction during the swing phase. It is to avoid the swing leg colliding with the stance leg that also cost metabolically [30]. Thus, there must be a lower limit for the step width reduction with the walking speed that we did not detect within our trials’ ranges.

On the other hand, there are metabolic expenditures that COM may not explain [20]. For instance, the muscle coactivation associated with age [31] or uneven walking [18] are not reflected in the COM work. Step variabilities perhaps can be categorized similarly. The elevated variabilities indicate the corrective actions done to regulate every single step. It is shown that the 65% step variability increase coincides with the 5.9% elevated metabolic expenditure [23]. We can associate the COM lateral work variability partially with the step width variability. Therefore, assigning step width variability cost to the lateral work might not be completely accurate. Nevertheless, it is the best metric that is currently available, since the quantification of step variability is challenging. The rise of variability with age separated the estimated metabolic cost of older adults from young adults (Figure 4C). The increased step variabilities with age might be related to their central nervous system (CNS) state [15].

The growth of the second compartment energetic magnitude (Workman model) with the terrain complexity also indicates that considerable effort and energy must be exerted to maintain the body balance and posture during uneven walking. The Workman model, despite its oversimplification, can provide some insight into the energetics of walking balance. Like COM lateral work, it shows that the balance cost is terrain-specific, and restricted lookahead and age contribute to the cost increase. However, comparing its cost with the COM lateral work proposes a gross difference. Thus, in addition to variability, the Workman model must also include the other costs that are not explained by the work. Over uneven terrains, preserving the posture or walking with flexed knees [18] may introduce further metabolic costs [32] that are reflected in the Workman model.

In conclusion, we have shown that the lateral mechanical work increases with the terrain amplitude and restricted lookahead to maintain the lateral balance. The elevated lateral work must have stemmed from increased step width and transition impulses that we might associate with sub-optimal foot landings. Nevertheless, we might have overestimated the associated metabolic costs since elasticity role in step work performances are recognized [18,22]. We could also account for the COM lateral work variabilities as portion of factors that the COM work does not reflect their energetic costs. Finally, although Workman model and our estimation depicted that lateral costs are terrain specific, grossly large Workman model estimate must encompass much more metabolic costs that cannot be reflected in the COM lateral work.

## Acknowledgements

This work was supported in part by the Natural Sciences and Engineering Research Council of Canada (NSERC) Discovery and Canada Research Chair (Tier 1) programs.

## References

[1] C.E. Bauby, A.D. Kuo, Active control of lateral balance in human walking, J Biomech 33 (2000) 1433–1440. 10.1016/s0021-9290(00)00101-9.

[2] M. Garcia, A. Chatterjee, A. Ruina, M. Coleman, The simplest walking model: stability, complexity, and scaling, J Biomech Eng 120 (1998) 281–288. 10.1115/1.2798313.

[3] S.M. Bruijn, J.H. Van Dieën, Control of human gait stability through foot placement, J. R. Soc. Interface. 15 (2018) 20170816. 10.1098/rsif.2017.0816.

[4] P.G. Adamczyk, A.D. Kuo, Redirection of center-of-mass velocity during the step-to-step transition of human walking, J Exp Biol 212 (2009) 2668–2678. 10.1242/jeb.027581.

[5] A.D. Kuo, Energetics of actively powered locomotion using the simplest walking model, J Biomech Eng 124 (2002) 113–120. 10.1115/1.1427703.

[6] J.M. Donelan, R. Kram, A.D. Kuo, Mechanical work for step-to-step transitions is a major determinant of the metabolic cost of human walking, Journal of Experimental Biology 205 (2002) 3717–3727. 10.1242/jeb.205.23.3717.

[7] J. Maxwell Donelan, R. Kram, K. Arthur D., Mechanical and metabolic determinants of the preferred step width in human walking, Proc. R. Soc. Lond. B 268 (2001) 1985–1992. 10.1098/rspb.2001.1761.

[8] S.M. O’Connor, H.Z. Xu, A.D. Kuo, Energetic cost of walking with increased step variability, Gait Posture 36 (2012) 102–107. 10.1016/j.gaitpost.2012.01.014.

[9] J.S. Matthis, J.L. Yates, M.M. Hayhoe, Gaze and the Control of Foot Placement When Walking in Natural Terrain, Curr Biol 28 (2018) 1224-1233.e5. 10.1016/j.cub.2018.03.008.

[10] J.E.A. Bertram, Constrained optimization in human walking: cost minimization and gait plasticity, Journal of Experimental Biology 208 (2005) 979–991. 10.1242/jeb.01498.

[11] A.D. Kuo, A simple model of bipedal walking predicts the preferred speed-step length relationship, J Biomech Eng 123 (2001) 264–269. 10.1115/1.1372322.

[12] O. Darici, H. Temeltas, A.D. Kuo, Optimal regulation of bipedal walking speed despite an unexpected bump in the road, PLoS ONE 13 (2018) e0204205. 10.1371/journal.pone.0204205.

[13] S.M. O’Connor, A.D. Kuo, Direction-Dependent Control of Balance During Walking and Standing, Journal of Neurophysiology 102 (2009) 1411–1419. 10.1152/jn.00131.2009.

[14] J.S. Matthis, B.R. Fajen, Humans exploit the biomechanics of bipedal gait during visually guided walking over complex terrain, Proc. R. Soc. B. 280 (2013) 20130700. 10.1098/rspb.2013.0700.

[15] F.A. Sorond, Y. Cruz-Almeida, D.J. Clark, A. Viswanathan, C.R. Scherzer, P. De Jager, A. Csiszar, P.J. Laurienti, J.M. Hausdor?, W.G. Chen, L. Ferrucci, C. Rosano, S.A. Studenski, S.E. Black, L.A. Lipsitz, Aging, the Central Nervous System, and Mobility in Older Adults: Neural Mechanisms of Mobility Impairment, J Gerontol A Biol Sci Med Sci 70 (2015) 1526–1532. 10.1093/gerona/glv130.

[16] A. Aboutorabi, M. Arazpour, M. Bahramizadeh, S.W. Hutchins, R. Fadayevatan, The effect of aging on gait parameters in able-bodied older subjects: a literature review, Aging Clin Exp Res 28 (2016) 393–405. 10.1007/s40520-015-0420-6.

[17] J.M. Workman, B.W. Armstrong, Metabolic cost of walking: equation and model, J Appl Physiol (1985) 61 (1986) 1369–1374. 10.1152/jappl.1986.61.4.1369.

[18] A.S. Voloshina, A.D. Kuo, M.A. Daley, D.P. Ferris, Biomechanics and energetics of walking on uneven terrain, Journal of Experimental Biology (2013) jeb.081711. 10.1242/jeb.081711.

[19] A.S. Voloshina, A.D. Kuo, D.P. Ferris, C.D. Remy, A model-based analysis of the mechanical cost of walking on uneven terrain, Bioengineering, 2020. 10.1101/2020.06.15.152330.

[20] J.M. Donelan, R. Kram, A.D. Kuo, Mechanical work for step-to-step transitions is a major determinant of the metabolic cost of human walking, Journal of Experimental Biology 205 (2002) 3717–3727. 10.1242/jeb.205.23.3717.

[21] A.D. Kuo, J.M. Donelan, A. Ruina, Energetic consequences of walking like an inverted pendulum: step-to-step transitions, Exerc Sport Sci Rev 33 (2005) 88–97. 10.1097/00003677-200504000-00006.

[22] G.A. Cavagna, N.C. Heglund, C.R. Taylor, Mechanical work in terrestrial locomotion: two basic mechanisms for minimizing energy expenditure, American Journal of Physiology-Regulatory, Integrative and Comparative Physiology 233 (1977) R243–R261. 10.1152/ajpregu.1977.233.5.R243.

[23] S.M. O’Connor, H.Z. Xu, A.D. Kuo, Energetic cost of walking with increased step variability, Gait & Posture 36 (2012) 102–107. 10.1016/j.gaitpost.2012.01.014.

[24] S. Gazzellini, M.L. Lispi, E. Castelli, A. Trombetti, S. Carniel, G. Vasco, A. Napolitano, M. Petrarca, The impact of vision on the dynamic characteristics of the gait: strategies in children with blindness, Exp Brain Res 234 (2016) 2619–2627. 10.1007/s00221-016-4666-9.

[25] Z.R. Kahaki, A.R. Safarpour, H. Daneshmandi, The spatiotemporal gait parameters among people with visual impairment: A literature review study, Oman J Ophthalmol 16 (2023) 427–433. 10.4103/ojo.ojo_24_23.

[26] M.G. Browne, J.R. Franz, Does dynamic stability govern propulsive force generation in human walking?, R. Soc. Open Sci. 4 (2017) 171673. 10.1098/rsos.171673.

[27] H. Nagano, L. James, W.A. Sparrow, R.K. Begg, Effects of walking-induced fatigue on gait function and tripping risks in older adults, J NeuroEngineering Rehabil 11 (2014) 155. 10.1186/1743-0003-11-155.

[28] N. Lythgo, R. Begg, R. Best, Stepping responses made by elderly and young female adults to approach and accommodate known surface height changes, Gait Posture 26 (2007) 82–89. 10.1016/j.gaitpost.2006.07.006.

[29] M.J. Major, R.L. Stine, S.A. Gard, The effects of walking speed and prosthetic ankle adapters on upper extremity dynamics and stability-related parameters in bilateral transtibial amputee gait, Gait & Posture 38 (2013) 858–863. 10.1016/j.gaitpost.2013.04.012.

[30] K.A. Shorter, A. Wu, A.D. Kuo, The high cost of swing leg circumduction during human walking, Gait & Posture 54 (2017) 265–270. 10.1016/j.gaitpost.2017.03.021.

[31] D.S. Peterson, P.E. Martin, Effects of age and walking speed on coactivation and cost of walking in healthy adults, Gait Posture 31 (2010) 355–359. 10.1016/j.gaitpost.2009.12.005.

[32] K.E. Gordon, D.P. Ferris, A.D. Kuo, Metabolic and mechanical energy costs of reducing vertical center of mass movement during gait, Arch Phys Med Rehabil 90 (2009) 136–144. 10.1016/j.apmr.2008.07.014.

